# Vimentin plays a functional role in mammary gland regeneration

**DOI:** 10.1101/134544

**Authors:** Reetta Virtakoivu, Emilia Peuhu, Anja Mai, Anni Wärri, Johanna Ivaska

## Abstract

In the mammary gland, vimentin intermediate filaments are expressed in stromal cells and in basal epithelial cell populations including gland-reconstituting mammary stem cells (MaSC), with largely undefined functions. Here, we studied how vimentin deficiency affects mouse mammary gland development. Our results demonstrate that in adult vimentin knockout mice (*Vim-/-*) mammary ductal outgrowth is delayed. The adult *Vim-/-* glands are characterised by dilated ducts, an imbalance in the proportion of basal to luminal mammary epithelial cells and a reduction in cells expressing Slug (Snai2), an established MaSC regulator. All of these features are indicative of reduced progenitor cell activity. Accordingly, isolated *Vim-/-* mammary epithelial cells display reduced capacity to form mammospheres, and altered organoid structure, compared to wt counterparts, when plated in a 3D matrix *in vitro*. Importantly, altered basal epithelial cell number translates into defects in *Vim-/-* mammary gland regeneration *in vivo* in cleared fat pad transplantation studies. Furthermore, we show that vimentin contributes to stem-like cell properties in triple negative MDA-MB-231 breast cancer cells, wherein vimentin depletion reduces tumorsphere formation and alters expression of breast cancer stem cell-associated surface markers. Together, our findings identify vimentin as a positive regulator of stemness in the developing mouse mammary gland and in breast cancer cells.

## Introduction

The mammary gland is a highly dynamic organ that develops through branching morphogenesis in puberty, evolves during the menstrual cycle, and undergoes terminal differentiation/dedifferentiation during pregnancy, lactation and involution. Increasing evidence suggests that the mammary epithelium in both humans and mice comprises a hierarchy of cells, spanning from bipotent mammary gland stem cells (MaSCs) to differentiated luminal and basal epithelial cells (Rios et al, 2014, Van Keymeulen et al, 2011, Visvader & Stingl, 2014). The MaSCs and the basal and luminal progenitor cells are considered particularly important for ductal elongation during pubertal growth and for lobulo-alveolar expansion during pregnancy (Tiede & Kang, 2011). Several marker proteins have been indicated for stem and progenitor cells in the mammary epithelium (Visvader & Stingl, 2014). The current view suggests that the transcription factors Sox9 and Slug (Snai2) (Guo et al, 2012), several integrin adhesion receptors (Rangel et al, 2016, Taddei et al, 2008), and molecules involved in Notch and Wnt signalling pathways (Bouras et al, 2008, Zeng & Nusse, 2010) are expressed in the MaSCs and committed progenitor cells that localize to the basal mammary epithelial niche (Shackleton et al, 2006).

Vimentin is a cytoskeletal type III intermediate filament protein widely used as a marker for mesenchymal cells (Coulombe & Wong, 2004). Despite the widespread expression of vimentin, the phenotype of vimentin knockout mice (*Vim*^*tm1Cba*^, here after called *Vim-/-*) is mild (Colucci-Guyon et al, 1994). Nevertheless, defects in motor coordination (Colucci-Guyon et al, 1999), wound healing (Cheng et al, 2016, Eckes et al, 2000) and endothelial function (Nieminen et al, 2006) accompany vimentin loss in these animals. Slug regulates the expression of vimentin (Vuoriluoto et al, 2011) alongside other genes involved in epithelial-to-mesenchymal transition (EMT), a key developmental program often activated during cancer invasion and metastasis (Mani et al, 2008). Interestingly, in vimentin-deficient cells, also Slug levels are substantially reduced, suggesting the existence of mutual regulatory pathways that control Slug and vimentin expression (Cheng et al, 2016, Virtakoivu et al, 2015). Vimentin has been shown to promote extracellular signal-regulated kinase (ERK)-mediated Slug phosphorylation, Slug-dependent vimentin and receptor tyrosine kinase Axl expression and MDA-MB-231 breast cancer cell invasion (Virtakoivu et al, 2015).

Other intermediate filament proteins such as nestin and GFAP have been implicated in self-renewal of neural stem cells (Park et al, 2010). Although Slug and vimentin are both expressed in basal mouse mammary epithelial cells (bMMECs), including the normal tissue reconstituting MaSCs (Soady et al, 2015, Ye et al, 2015), the functional role of vimentin in mammary gland development and stemness has not been previously studied. Here, we report that *Vim-/-* mouse mammary epithelium has delayed ductal outgrowth and reduced regenerative capacity *in vivo* and demonstrate that vimentin is an important regulator of self-renewal capacity in normal and transformed mammary epithelial cells.

## Results and Discussion

### Vimentin is expressed in basal epithelial and stromal cells in the mouse mammary gland

Supporting previous observations (Soady et al, 2015, Ye et al, 2015), vimentin was found to be expressed in basal mammary epithelial cells, and in stromal cells in human and mouse mammary gland tissue sections (Fig. A, B). Vimentin expression was lower in the epithelial compartment than in the stroma although some basal cells particularly in the mouse mammary gland exhibited higher vimentin expression level (Fig. 1B). In addition, quantitative PCR analysis of vimentin mRNA expression in mouse mammary gland cell populations (basal epithelial cells, luminal progenitor and mature luminal epithelial cells and stromal cells, sorted by flow cytometry based on CD24 and ICAM1 surface expression (Di-Cicco et al, 2015) revealed notable vimentin levels in stromal cells and to a lesser extent in basal epithelial cells (Fig. 1C). As expected, luminal progenitor cells and mature luminal epithelial cells did not express vimentin (Fig. 1C).

**Figure 1.**
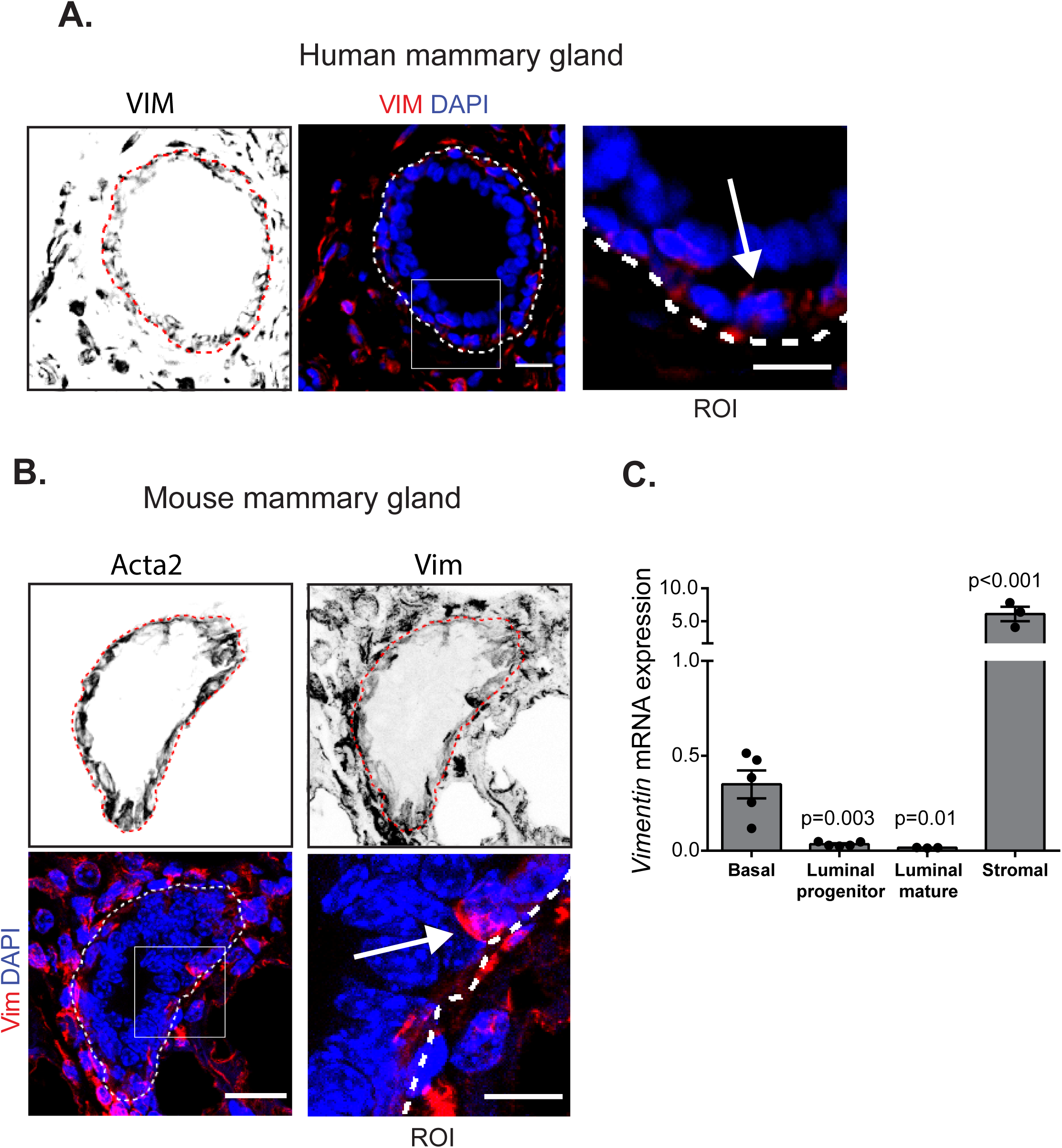
Vimentin is expressed in basal epithelial and stromal cells in the mammary gland. **A-B**. Immunohistochemical (IHC) analysis of vimentin expression in human (**A**) and mouse mammary glands (**B**). Cross-sections of vimentin- and DAPI-labelled (**A**; single image section) or vimentin-, Acta2 (basal cells) and DAPI-labelled (**B**; maximum projection image) mammary ducts and magnification of the region of interest (ROI) are shown. The approximate position of the basement membrane (dashed line) and vimentin-positive basal cells in the ROI (arrow) are indicated. Scale bars represent 20 μm (original image) and 10 μm (ROI). **C.** Quantitative PCR analysis of vimentin mRNA expression in mouse mammary gland cell populations sorted by FACS into basal epithelial cells (Lin^neg^CD24^int^ICAM-1^hi^), luminal progenitor cells (Lin^neg^CD24^hi^ICAM-1^int^), mature luminal epithelial cells (Lin^neg^CD24^hi^ICAM-1^neg^), and stromal cells (Lin^neg^CD24^neg^) (mean ± SEM, n = 3 − 5). Statistical analysis, unpaired Student’s t-test.

### Mammary ductal outgrowth is delayed in vimentin knockout mouse

Previous reports describing vimentin expression in the mouse MaSC compartment (Soady et al, 2015, Wang et al, 2015, Ye et al, 2015) led us to investigate the *Vim-/-* and wt control mouse mammary glands. Lack of vimentin expression in the *Vim-/-* mammary gland was confirmed by IHC (Fig 2A). In laboratory mice, sexual maturity and mammary gland maturation is reached by 8 weeks after birth (Green & Witham, 2009). Mammary gland whole mount analyses, performed at the same phase of the oestrus cycle, revealed impaired mammary ductal outgrowth in adult virgin *Vim-/-* mice (Fig. 2B-E). The stunted growth of the mammary ductal network, clearly visible in 10-week-old female *Vim-/-* mice (Fig. 2B,D), was no longer apparent in older animals by 15 weeks of age (Fig. 2C,E). Thus, these data demonstrate that mammary ductal outgrowth is significantly delayed but not terminally impaired in the absence of vimentin.

**Figure 2.**
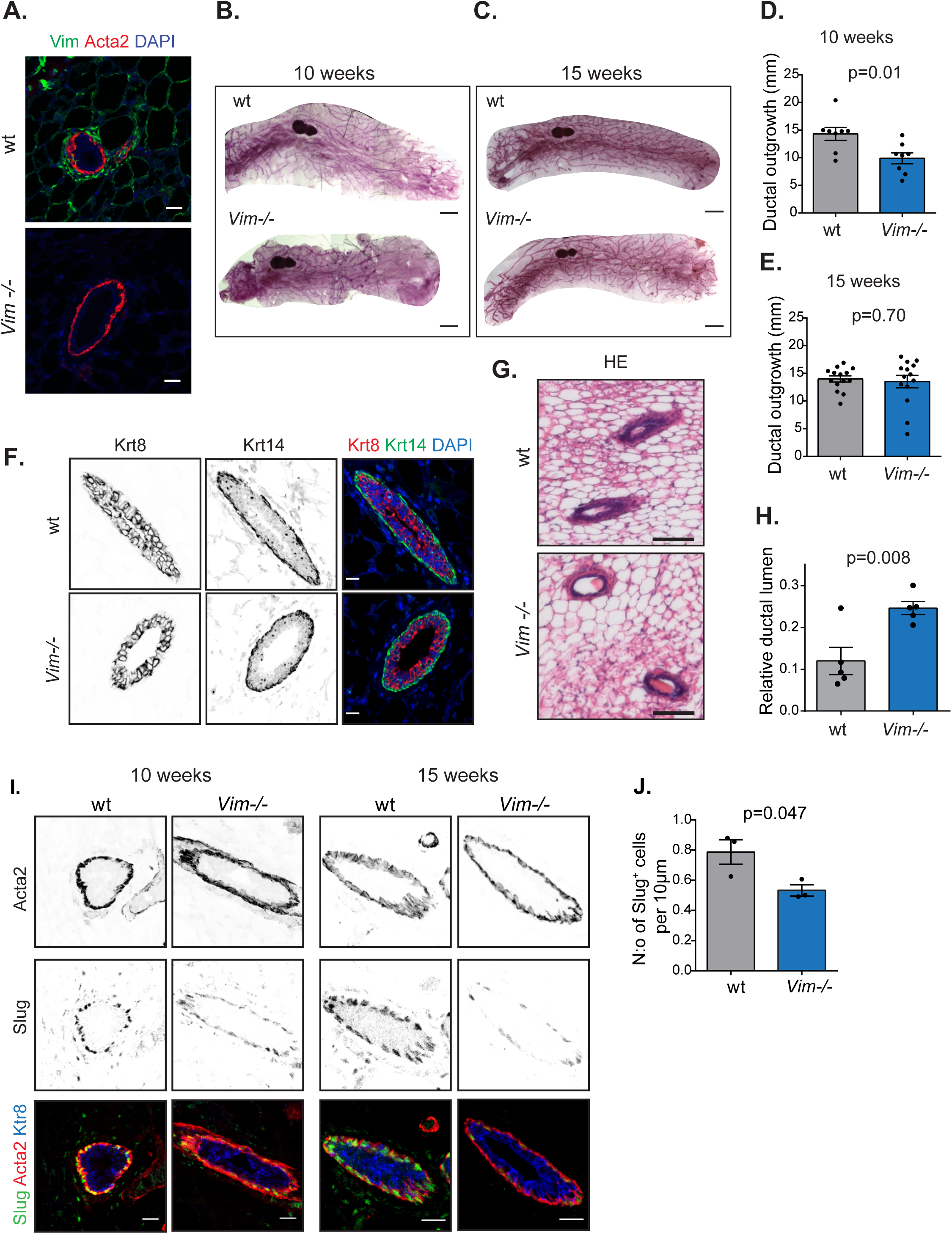
Mammary ductal outgrowth is delayed in vimentin knockout mice. **A.** Representative mammary gland tissue sections from 15-week-old wt and *Vim-/-* female mice were labelled for nuclei (DAPI), vimentin and Acta2 by IHC-P (n = 3 mice). Scale bar, 10 μm. **B-C.** Representative images of mammary gland whole mounts from 10 weeks (**B**) and 15 weeks (**C**) old wt and *Vim-/-* mice. Scale bar 2 mm. **D-E.** Quantification of mammary ductal outgrowth beyond the inguinal lymph node in mammary gland whole mounts from 10 weeks (**D.**) and 15 weeks (**E.**) old wt and *Vim-/-* mice. (n_10 weeks_ = 8 mice, n_15 weeks_ = 12 mice) (mean ± SEM). Statistical analysis, unpaired Student’s t-test. **F.** Mammary gland tissue sections from 15 weeks old wt and *Vim-/-* female mice were labelled for keratin-8 (Krt8; Luminal epithelial) and keratin-14 (Krt14; Basal epithelial) expression by IHC-P (n=5−6 mice). Scale bar 10 μm. **G-H**. Hematoxylin-eosin (HE) stained mammary gland tissue sections from 15 weeks old wt and *Vim-/-* female mice (**G.**) and quantification of mammary ductal lumen area (**H.**). (n = 5 mice; 20-22 ducts per mouse). Scale bar, 100 μm. Mean ± SEM. Unpaired Student’s t-test. **I-J.** Mammary gland tissue sections from 10 weeks old (left) and 15 weeks old (right) wt and *Vim-/-* female mice were labelled for Acta2, Slug and Keratin 8 (krt8) by IHC-P (**I.**). Scale bar 20 μm. The number of Slug positive cells per 10 μm distance of basement membrane was quantified from 15 weeks old mice (**J.**). (Mean ± SEM; n=3 mice; 10 ducts analysed per animal). Unpaired Student’s t-test.

Establishment of correct polarity is critical for normal mammary gland development and is regulated by the co-ordinated actions of adhesion receptors and the cell cytoskeleton. Given that vimentin interacts with integrins and regulates focal adhesion turnover (Kim et al, 2016, Kreis et al, 2005, Liu et al, 2015), we wanted to investigate whether the lagging mammary gland outgrowth is linked to perturbed organization of the mammary bilayer in the *Vim-/-* mice. Although normal mammary epithelial polarisation was observed in adult *Vim-/-* mammary ducts based on the distribution of established markers: keratin-8 (Krt8; luminal) and keratin-14 (Krt14; basal) (Fig. 2F), the lumens of these ducts appeared to be larger compared to those of age-matched 15-week-old wt mice (Fig. 2G-H). For comparable quantification, the measurements were restricted to the relative area of the lumen in perpendicular cross sections of the smaller ducts at corresponding areas (Fig. 2H).The enlarged lumen in *Vim-/-* mouse mammary ducts resembles a benign breast condition called ductal ectasia, the dilation of mammary ducts (Rahal et al, 2011) that in humans and mice is related to aging. In normal mice, ductal ectasia occurs at about two years of age, when progenitor activity is already reduced (Jackson et al, 2015). Increased levels of mammary stem/progenitor cells inhibit ectasia in aged mice (Jackson et al, 2015) suggesting that the accelerated occurrence of ectasia in the *Vim-/-* mice could also be related to reduced self-renewal capacity and MaSC content.

The bMMEC layer where vimentin expression was detected (Fig. 1A-C) is also the niche for the MaSC population (Shackleton et al, 2006), and vimentin is a well-characterized target gene of Slug (Vuoriluoto et al, 2011) that is expressed in bipotent MaSCs (Guo et al, 2012). Since reduced Slug levels have previously been detected in vimentin-deficient cells (Cheng et al, 2016, Virtakoivu et al, 2015), the expression of Slug was investigated in adult (10 or 15 weeks old) *Vim-/-* and wt mouse mammary gland tissue sections by IHC (Fig. 2J). In the wt mammary gland Slug labelling resulted in a previously reported basal epithelial expression pattern (Guo et al, 2012) (Fig, 2J). While the *Vim-/-* mammary epithelium also contained Slug-positive basal cells (Fig. 2J), the amount of these cells was significantly reduced (Fig. 2K). These data suggest that vimentin could regulate the basal MaSC/progenitor cell population in the mouse mammary gland.

### Vimentin deficiency leads to reduced proportion of basal mammary epithelial cells

To evaluate the basal to luminal MMEC ratio, we isolated MMECs from 15-week-old wt and *Vim-/-* mice, and surface labelled single cell suspensions to detect CD24 expression in combination with CD29 (integrin beta 1) or CD49f (integrin alpha 6). From lineage negative cells (CD31^neg^CD45^neg^), the proportion of basal (CD24^int^CD29^+^, CD24^int^CD49f^+^) and luminal (CD24^hi^CD29^neg^, CD24^hi^CD49f^neg^) MMECs was then quantified by flow cytometry (Fig. 3A-B) (Taddei et al, 2008). Vimentin deficiency led to significantly (p<0.05) or almost significantly (p=0.05) reduced proportion of bMMECs with both the employed labelling strategies (Fig. 3A-B) as well as with an alternative basal/mature luminal epithelial labelling against CD61 (integrin beta3) in combination with CD29 and CD49f (Fig. S1A-B), suggesting that the mammary epithelial compartment harbouring the MaSC/progenitor cells is diminished in the absence of vimentin.

**Figure 3.**
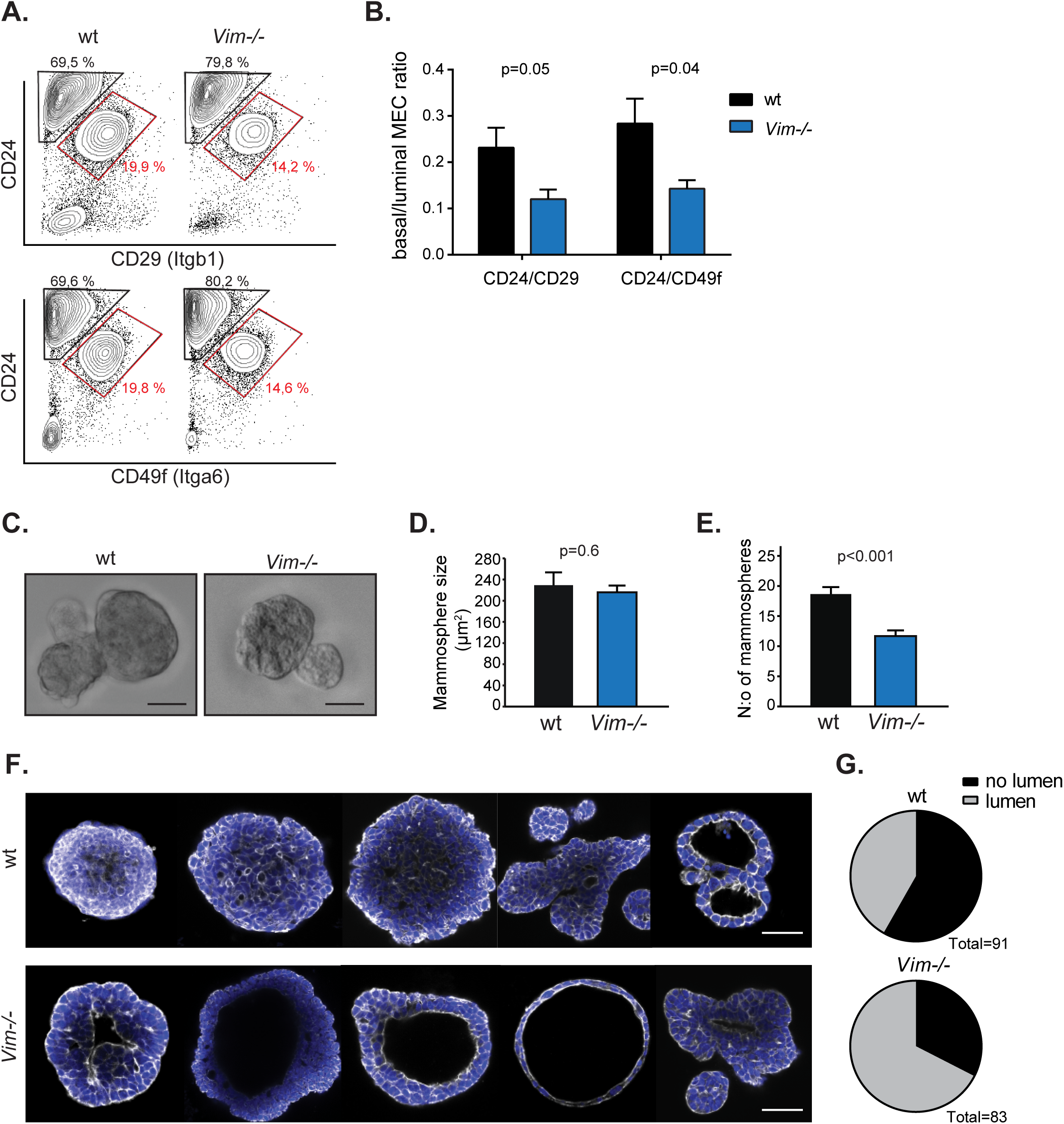
Vimentin knockout MMECs contain reduced proportion of basal cells, and form less mammospheres and filled organoids *in vitro*. **A-B.** MMECs were isolated from 15-18 weeks old wt and *Vim-/-* female mice and immunolabelled for surface markers. From the lineage negative cells (CD31^neg^CD45^neg^), the basal epithelial (CD24^int^CD29^+^, CD24^int^CD49f^+^; gate and % of cells indicated in red) and luminal epithelial (CD24^hi^CD29^neg^, CD24^hi^CD49f^neg^) cell populations were quantified by flow cytometry (**A.**). The ratio between basal and luminal epithelial cells in each sample was calculated (**B.**). n= 3-4; Mean ± SEM. Unpaired Student’s t-test. **C-E.** Wt and *Vim-/*-MMECs were cultured in low attachment conditions to examine the capacity to form mammospheres. Mammospheres were imaged by bright field microscopy (**C.**) and the size (**D.**) and number (**E.**) of mammospheres per well (total 6 wells/ genotype) was quantified. Representative result of two independent experiments is shown. Scale bar, 50 μm. Mean ± SEM. Unpaired Student’s t-test. **F-G.** The middle plain of organoids formed by wt and *Vim-/*-MMECs in 3D Matrigel(tm) cultures were imaged by confocal microscopy. Representative images of distinct lumen morphologies observed are shown (**F.**) and the fraction of organoids with or without lumen was quantified (**G.**). Data was pooled from three independent experiments, each 20-40 organoids per experiment. Scale bar, 50 μm. Fisher’s exact test (p=0.0008).

### MMECs from vimentin knockout mice form less mammospheres and filled organoids

Mammary progenitor cells can be distinguished based on their ability to generate and propagate colonies in suspension *in vitro*. Both human and mouse MaSCs can form mammospheres in suspension (Dontu et al., 2003; Liao et al., 2007) and the ability of cells to generate mammospheres reflects the number of self-renewing, regenerative MaSCs within the cell population (Liao et al., 2007). To investigate the ability of *Vim-/-* and wt MMECs to grow as mammospheres, equal numbers of *Vim-/-* and wt MMECs were seeded at low density in low attachment conditions. While the size of *Vim-/-* mammospheres was comparable to wt, the number of mammospheres formed by *Vim-/-* MMECs was significantly lower (Fig. 3C-E). These data demonstrate that primary *Vim-/*-MMECs have a reduced level of self-renewing cells compared to their wt counterparts. Similarly, MMECs isolated from Slug knockout mice have reduced ability to generate mammospheres *in vitro* (Nassour et al, 2012), suggesting that vimentin and Slug could jointly contribute to a common self-renewal pathway in mammary epithelial cells.

MMECs cultured in laminin-rich reconstituted basement membrane (rBM) have similar features to mammary epithelium *in vivo*, including the formation of acini-like organoids with a hollow lumen and apico-basal polarity. The mammary ductal lumen is formed when the inner cell population is cleared by anoikis-like cell death mechanisms (Mailleux et al, 2007). Interestingly, *Vim-/-* MMECs that were seeded as single cells in three dimensional (3D) rBM formed organoids with altered morphology compared to wt MMECs. Vimentin-deficient organoids formed a lumen significantly more often than the wt organoids (Fig. 3F-G). This observation could be related to the reduced proportion of basal epithelial cells in the *Vim-/-* mammary gland as the luminal population predominantly forms hollow acini in traditional Matrigel colony-forming assays while the basal cell population tends to form a more heterogeneous array of structures, including ductal forms and solid spheroids (Shackleton et al., 2006; Stingl et al., 2006). Interestingly, this *in vitro* phenotype resembles the enlarged mammary epithelial lumen in *Vim-/-* tissue sections *in vivo* (Fig. 2H-I). Together, our data suggests that vimentin expression regulates basal cell compartment and the number of self-renewing MaSCs in the mouse mammary epithelium.

### Vimentin silencing reduces tumorsphere formation in MDA-MB-231 breast carcinoma cells

The existence and regulation of stem cell populations has been a matter of intense research in the context of breast cancer. Previously, vimentin expression was shown to increase during serial tumorsphere culture of several breast cancer cell lines (Borgna et al, 2012), and increased vimentin and Slug expression was demonstrated in a low adherent and more metastatic side population of MDA-MB-231 human breast cancer cells (Morata-Tarifa 2016). To investigate whether vimentin regulates the stem cell associated features of the triple-negative and mesenchymal-like MDA-MB-231 breast cancer cells, vimentin was silenced with an siRNA smartpool (Fig. 4A), previously reported to be specific for vimentin with no evidence of off-target effects using gold-standard controls (Virtakoivu et al, 2015). CD49f and CD61, in particular, have been described as markers of breast cancer stem cells (Brooks et al, 2016, Vaillant et al, 2008). Interestingly, vimentin knockdown resulted in a significant downregulation of CD49f and CD61 cell surface expression (Fig. 4B). Importantly, in low attachment assays, MDA-MB-231 tumorsphere formation was reduced in the absence of vimentin based on the number of spheres (Fig. 4D) and on total protein content of spheres generated by equal numbers of seeded vimentin-silenced or control cells (Fig. 4E). These results indicate that vimentin influences tumorsphere formation capacity and the expression of integrin adhesion receptors in breast cancer cells *in vitro*. In breast carcinomas, vimentin expression is most prevalent in the triple negative subtype (61% vimentin positive) and vimentin positivity correlates strongly with nuclear Slug expression (Virtakoivu et al, 2015). Accordingly, vimentin expression is typically associated with high tumour invasiveness and resistance to chemotherapy (Korsching et al, 2005, Yamashita et al, 2013). It has also been suggested that vimentin-expressing breast carcinomas could be derived from the bipotent breast progenitor cells (Korsching et al, 2005). Our data demonstrating that vimentin influences tumorsphere formation in breast cancer cells supports this hypothesis. Recently, vimentin knockdown in orthotopic breast cancer allografts in a hyperinsulinemia mouse model was shown to reduce pulmonary metastasis (Zelenko et al, 2017). Together, these findings suggest that vimentin contributes to breast cancer initiation and progression.

**Figure 4.**
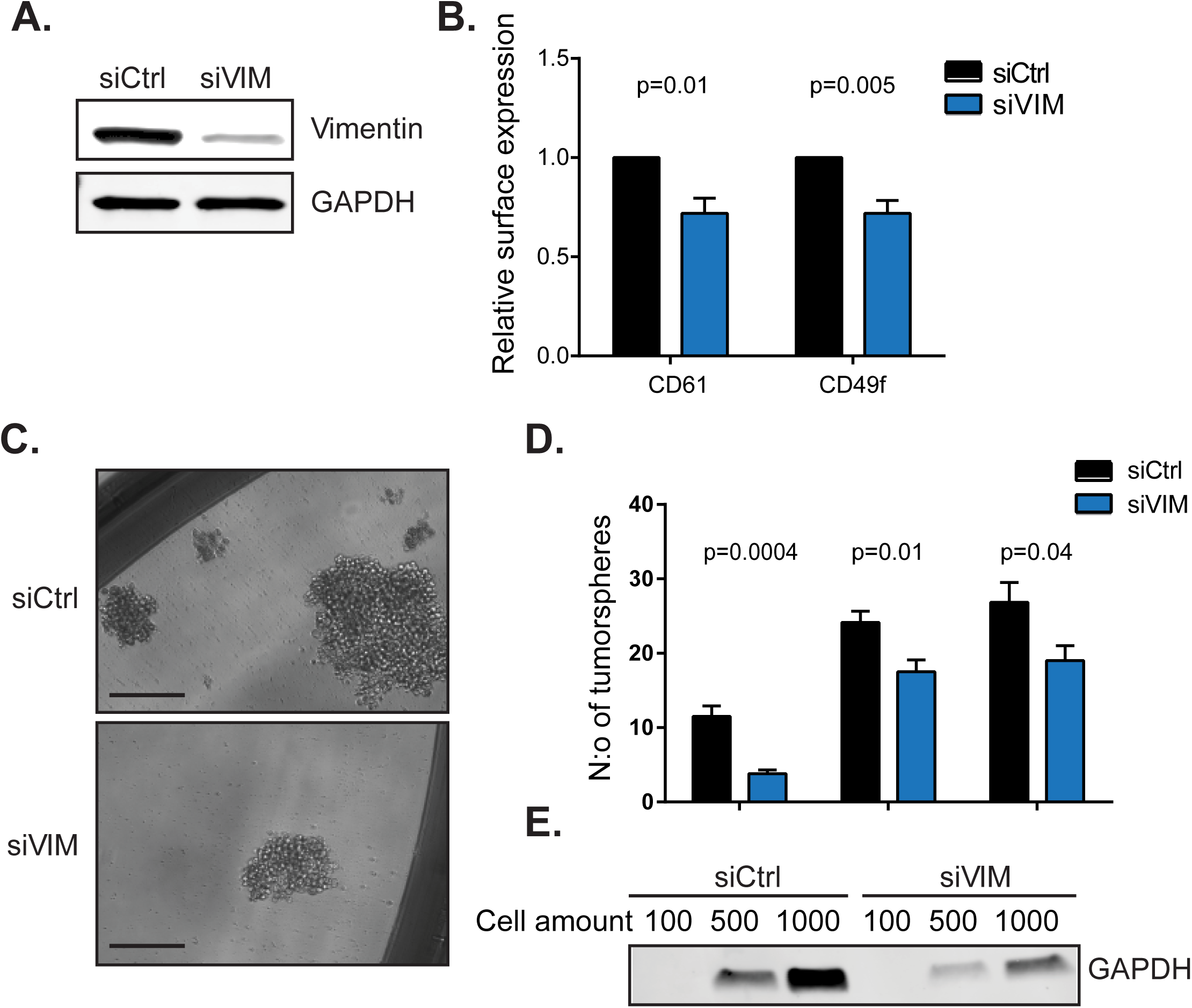
Vimentin silencing reduces tumorsphere formation in MDA-MB-231 human breast carcinoma cells. **A.** Silencing of vimentin expression in MDA-MB-231 breast cancer cells by siRNA smartpool was verified by western blotting from cell lysates. Control cells were treated with control siRNA. **B.** Relative expression of cell surface integrin adhesion receptors in control or vimentin silenced MDA-MB-231 cells was analysed by flow cytometry (n=6−7 from minimum 4 experiments). Mean ± SEM. Paired t-test. **C-E.** Formation of tumorspheres by control or vimentin silenced MDA-MB-231 cells. Tumorspheres were imaged with bright field microscopy (C.) (Scale bar, 400 μm), and their number scored per well (D.). The protein content of tumorspheres generated by equal numbers of vimentin silenced and control cells was analysed by loading equal volumes of cell lysate in SDS-PAGE gels and blotting for the expression of house-keeping protein GAPDH (E.). Representative results of 3 independent experiments are shown. Mean ± SEM. Unpaired Student’s t-test.

### Regenerative capacity is reduced in vimentin knockout mouse mammary glands

Finally, to evaluate how vimentin affects MaSC capacity *in vivo*, we grafted adult wt or *Vim-/-* mammary gland pieces into epithelium-free fat pads of prepubertal wt recipient mice. Vimentin deficiency did not influence the outgrowth from primary transplants when the growth area was quantified 10 weeks post-transplantation (Fig. 5A-B). These data could not be directly compared to differential rates of normal mammary gland development in wt and *Vim-/-* mice (Fig. 2D-G) as the outgrowth from the transplants was much slower. Interestingly, when the primary transplants with noticeable epithelial growth (Fig. S1C) were further cut into pieces and re-transplanted in secondary recipients, *Vim-/-* transplants demonstrated a substantial loss in growth capacity 15 weeks post-transplantation (Fig. 5C-D). Additionally, the take-on-rate demonstrating the fraction of transplants that initiated growth was only compromised at the second round of transplantation (Fig. 5E). The inability to re-establish a ductal tree only in the secondary transplants strongly indicates a lack of functional stem cells in *Vim-/-* mammary epithelium. A similar defect in mammary gland self-renewal capacity was previously observed in mice where *Itgb1* was conditionally deleted in the basal epithelial cells (Taddei et al, 2008). The data presented here suggest that vimentin plays a functional role in the regulation of regenerative capacity in the mammary gland *in vivo*.

**Figure 5.**
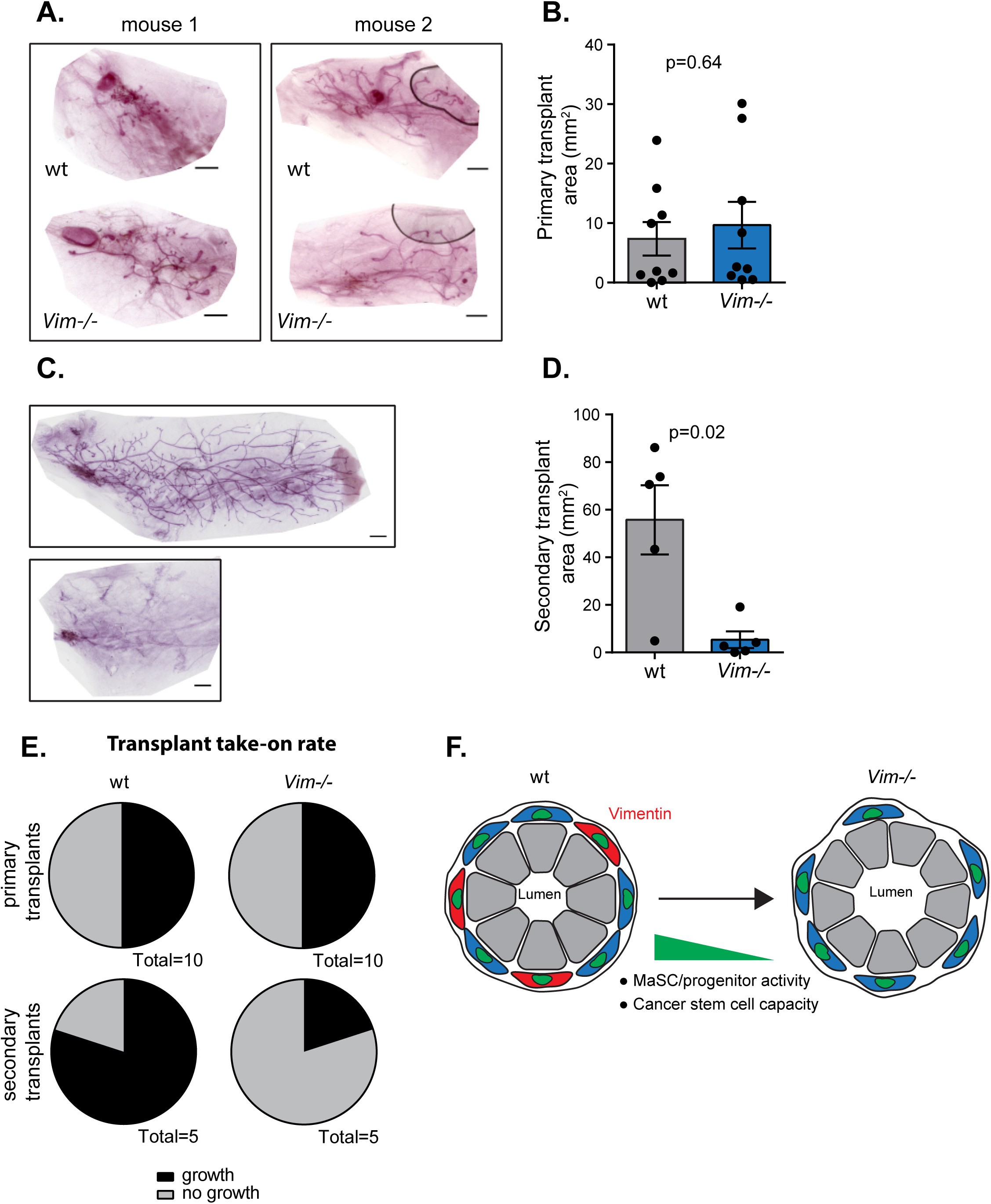
Regenerative capacity is reduced in vimentin knockout mouse mammary gland. **A-B.** Primary mammary gland transplantation. Mammary gland pieces from wt and *Vim-/-* donor mice were transplanted in cleared fat pads of prepubertal wt recipient mice. Mammary gland whole mounts were prepared 10 weeks later (A.) and the growth area of the transplant was quantified (B.). Two example whole mounts are shown. (n=10; Mean ± SEM). Unpaired Student’s t-test. **C-D.** Secondary mammary gland transplantation. Mammary gland pieces from wt and *Vim-/-* primary transplants with epithelial growth were further transplanted in cleared fat pads of prepubertal wt recipient mice. Mammary gland wholemounts were prepared 15 weeks later (**C.**) and the growth area of the secondary transplant was quantified (**D**.). n=5 Mean ± SEM. Unpaired Student’s t-test. **E.** Growth take-on-rate in primary (**A-B.**) and secondary (**C-D**.) mammary gland transplantation experiments with wt or *Vim-/-* mammary gland tissue pieces. **F.** Schematic model for the function of vimentin in regulation of mammary stem/progenitor cell activity and breast cancer stem cell capacity. Reduced proportion of basal epithelial cells (blue) and enlarged lumen size in vimentin-deficient mammary ducts are depicted. Vimentin expressing cells are labelled with red.

Our data demonstrate that vimentin regulates the rate of mammary ductal outgrowth, a process involving MaSCs, during mouse puberty, and affects the morphology of mature mammary ducts in vivo. Interestingly, Slug knockout mice also have delayed mammary gland development during puberty and MMECs isolated from these mice have reduced regenerative capacity of in vitro (Nassour et al, 2012), suggesting that vimentin and Slug regulate the same processes in the mammary epithelium. Together, our data suggest that Slug and vimentin mutually and positively regulate mammary gland stemness and potentially influence cancer stem cell properties in breast cancer cells. As a regulator of Slug expression, vimentin could even function upstream of Slug in the MaSC/progenitor compartment and thereby affect the regenerative capacity of the mammary gland.

## Conclusions

Here, we have investigated for the first time the functional consequences of vimentin loss for mammary gland biology. These studies were prompted by the established role of vimentin in EMT (Mendez et al, 2010), and the strong links between EMT-inducing factors and the self-renewing capacity of the mammary epithelium (Asiedu et al, 2014, Ye et al, 2015). We have earlier demonstrated that vimentin is required for EMT induced by Ras, Slug and TGFβ in cancer cells (Vuoriluoto et al, 2011). In addition, we have defined a signal-integrating function for vimentin on the ERK-MAPK-Slug-EMT axis where vimentin contributes to EMT progression (Virtakoivu et al, 2015). Here we identify vimentin as a regulator of regenerative capacity in the mouse mammary gland *in vivo* and *in vitro*, and demonstrate that vimentin also supports the formation of breast cancer tumorspheres indicative of a role in regulating cancer stem cell capacity (Fig. 5F). Importantly, vimentin appears to be a central node in stem cell programs operating in cancer and normal stem cells. Understanding how vimentin contributes to mammary stem cell regulation can be of importance in developing therapeutic strategies against breast cancer.

Furthermore, we demonstrate a delay in the development of *Vim-/-* mouse mammary ducts, characterised by the depletion of basal epithelial cells and the presence of enlarged lumens, which could be related to impaired mammary gland stem cell function (Fig. 5F). Interestingly, we find many parallels between the mammary gland defects in *Vim-/-* mice and those reported earlier for Slug knockout animals. The Slug knockout mouse has delayed mammary gland development during puberty and MMECs from these mice have reduced capacity to generate mammospheres (Nassour et al, 2012). In addition, other Slug-regulated mammary gland processes such as alveologenesis and involution (Castillo-Lluva et al, 2015, Desgrosellier et al, 2014) could be affected in the *Vim-/-* mice. The role of vimentin in the tertiary differentiation of mouse mammary gland will be studied in the future.

## Materials and methods

### Animals

Vimentin-deficient homozygote (*Vim*^*-/-*^) (Colucci-Guyon et al, 1999) and wt mice in mixed 129/Sv × C57BL/6 background were generated through heterozygote and homozygote mating. The PCR genotyping method was used to analyse the genotype of the mice. Age-matched female mice were used in all experiments. Mice were synchronized for estrus cycle in experiments where individual mice were examined. Otherwise, a minimum of two mice were pooled per MMEC isolation. For cell sorting and qPCR experiments, mammary glands were collected from adult virgin wt BALB/cByJ female mice. For frozen mammary gland sections, tissues were collected from adult virgin wt C57Bl/6 female mice. All animal studies were ethically performed and authorised by the National Animal Experiment Board and in accordance with The Finnish Act on Animal Experimentation (Animal licence numbers 7522/04.10.03/2012, ESAVI-5588-04.10.07-2014, ESAVI-9339-04.10.07-2016).

### Whole mount staining and analysis

The fourth mammary gland was dissected and placed on an object glass, left to adhere for few minutes, fixed in Carnoy’s medium (60% EtOH, 30% chloroform, 10% glacial acetic acid) overnight (o/n) at +4°C, followed by dehydration in decreasing EtOH series and staining with carmine alum (0.2% carmine, 0.5% aluminium potassium sulphate dodecahydrate) o/n at room temperature (RT). Samples were dehydrated and cleared in xylene for 2-3 days. Mounting was done in DPX Mountant for histology (Sigma) and images were taken with Zeiss SteREO Lumar V12 stereomicroscope (NeoLumar 0.8× objective, Zeiss AxioCam ICc3 colour camera). All images per gland were combined automatically into a mosaic picture with PhotoShop. Ductal outgrowth was analysed in ImageJ by measuring the distance of the ductal tree from the inguinal lymph node into the fat pad (adult mice) or by measuring the area covered by the ductal tree (transplantation).

### MMEC isolation

Ten weeks old or 15-18 weeks old female mice were sacrificed and the inguinal lymph node was removed. Tissues were minced with scalpel to small pieces and the homogenate was collected into a collagenase solution (DMEM/F12 media containing 2.5% FCS, 5 μg/mL insulin, 50 μg/mL gentamicin, and 200 mM glutamine and 20 mg of Collagenase XI (Sigma) for 1 g of tissue) and incubated at +37 °C with agitation (120 rpm) for 2-3 h. The resulting cell suspension was centrifuged at 1500 rpm and resuspended in DMEM/F12 medium (containing 1 % of penicillin/ streptomycin and 50 μg/mL gentamicin) and DNase I (20 U/ml) for 3 min. The sample was then subjected to five pulse centrifugation steps at 1500 rpm; every time the supernatant was carefully removed and the pellet was resuspended to 10 ml of DMEM/F12 medium. Next, organoids were incubated with Accutase (StemCell Technologies) to obtain a single cell suspension and cells were pipetted through a 70 μm cell strainer (BD Biosciences). Cells were used for mammosphere assays, flow cytometry and western blotting.

### Cell culture and transfections

MDA-MB-231 cells were cultured in DMEM supplemented with 10 % FBS, 1% L-glutamate and non-essential amino acids. Lipofectamine 3000 transfection reagent (Invitrogen) was used for siRNA transfections and the transfection was done according to the manufacturer’s instructions and as previously described (Virtakoivu et al, 2015). Silenced cells were used for experiments 48-72 h after transfection. Specific silencing of vimentin by single siRNA oligos in the vimentin siRNA Smartpool was previously validated by western blotting (Virtakoivu et al, 2015).

### Antibodies

The following antibodies were used in the study: For immunofluorescence and Western blotting, Acta2 (alpha smooth muscle actin; clone 1A4, Sigma), Krt8 (keratin 8, clone Troma-I, Hybridoma Bank), Krt14 (keratin 14, Covance), Slug (Snai2, C19G7, Cell Signalling), GAPDH (5G4 Mab 6C5, Hytest) and vimentin (VIM, V9, Santa Cruz; D21H3, Cell Signalling). For flow cytometry of mouse cells, CD45-Pacific Blue (clone 30-F11), CD31-Pacific Blue (clone 390), CD61-A647 (ITGB3, clone 2C9.G2), CD29-A488 (ITGB1, clone HMβ1-1), CD49f-A488 (ITGA6, clone GoH3) (all from BioLegend) and CD24-FITC (clone M1/69; BD Biosciences) antibodies were used. For flow cytometry of human cells, CD61 (ITGB3, MCA728, Serotec), and CD49f (ITGA6, MCA699, clone GoH3, Serotec) antibodies were used. AlexaFluor-conjugated (Life Technologies) and HRP-linked (GE Healthcare) secondary antibodies against rat, rabbit and mouse IgG were used in immunolabelling, flow cytometry and Western blotting.

### In vitro organoid and mammosphere assays

For organoid and mammosphere cultures, cells were counted and different amounts of cells were plated in 96-well low attachment plates (Corning) and allowed to proliferate for 6-9 days. For organoid assays the following medium was used; Epicult Base medium (StemCell Technologies) supplemented with Epicult supplements, 10 ng/ml EGF, 4 μg/ml heparin, 10 ng/ml bFGF, 5 μM Y-27632 ROCK inhibitor, 5 % matrigel and 5 % FCS. For mammosphere assays the same medium was used without ROCK inhibitor, Matrigel and FCS. When using vimentin or control siRNA silenced MDA-MB-231 cells for the mammosphere assays, the siRNA mix was added to the cells at day 3. Formed mammospheres were manually counted and images taken with EVOS Cell Imaging System (ThermoFisher Scientific). All wells / cell type / cell amount were pooled together and lysed for western blotting.

For immunofluorescence staining the organoids were gently collected and centrifuged at 700 rpm for 5 minutes followed by 4 % PFA fixation. Organoids were resuspended into 20 % matrigel in PBS and plated on matrigel coated 8-well Ibidi μ-slide wells (Ibidi GmbH). Next, organoids were permeabilized with 0.5 % Triton-X in PBS for 10 minutes, washed three times with 100 mM glycine in PBS and blocked with 5 % horse serum, 0.3 % triton-X, 0.2 % BSA, and 0.05% NaN_3_ in PBS for 1.5 h at RT. Phalloidin Atto-488 (1/50) (Sigma-Aldrich) was incubated o/n, organoids were washes three times with 0.3 % triton-X in PBS and nuclei stained with Hoecst (1/3000). Organoids were washed with PBS, mounted with Vectashield and left to dry for 1 h at + 37°C. Imaging was done with Carl Zeiss LSM780 laser-scanning confocal microscope and quantification with ImageJ software.

### Immunohistochemistry

Formalin-fixed paraffin-embedded human mammary gland tissues were collected from the archives of the Department of Pathology, Helsinki University Central Hospital, Helsinki, Finland. An Institutional Review Board of the Helsinki University Central Hospital approved the study. Mammary gland tissues destined for formalin or Carnoy’s medium fixation and paraffin embedding were collected from 10-15 weeks old female mice of the indicated genotype. Samples were deparaffinised and rehydrated. Epitope unmasking was performed in citrate buffer using 2100 Antigen retriever (Aptum, UK). Samples were blocked with 0.5% FCS in PBS for 45 min and incubated overnight in blocking buffer,followed by washing with PBS and incubation with fluorochrome-conjugated secondary antibodies for 2 h at RT. Samples were washed, stained with DAPI (4′,6-Diamidino-2-Phenylindole, Dihydrochloride; Life Technologies) and mounted in Mowiol containing DABCO^®^ (Sigma) anti-fading reagent. Imaging was conducted with a Zeiss Axiovert 200M inverted wide-field microscope (hematoxylin-eosin), 3i CSU-W1 spinning disk confocal microscope (Intelligent imaging innovations) with Hamamatsu CMOS Orca Flash 4 or with Carl Zeiss LSM780 laser scanning confocal microscope.

### Flow cytometry

Cells were suspended into Tyrodes buffer (10 mM HEPES-NaOH at pH 7.5, 137 mM NaCl, 2.68 mM KCl, 1.7 mM MgCl_2_, 11.9 mM NaHCO, 3.5 mM glucose, 0.1 % BSA) and approximately 5 x10^5^ cells were used per staining. In the case of MMEC cells, directly conjugated antibodies were added and incubated at +4°C for 30 minutes, washed twice and fixed with 4 % PFA. For staining the vimentin-silenced MDA-MB-231 cells, cells were first fixed with 4 % PFA, washed with PBS and suspended into PBS containing 1 % BSA. Primary antibodies were incubated for 30 minutes followed by washes and incubation with Alexa Fluor 488 /647 secondary antibodies (1/400). Samples were run with BD LSR Fortessa flow cytometer (BD Biosciences) and analysed with Flowing software (Cell Imaging Core, Turku Centre for Biotechnology).

### Cell preparation, sorting and qPCR

Samples were prepared as previously described (Di-Cicco et al, 2015, Peuhu et al. 2017). Single cells were prepared from inguinal mammary glands taken from virgin adult BALB/cByJ females according to a detailed protocol described previously (Di-Cicco et al, 2015). Freshly isolated cells were incubated at 4 °C for 20 min with the conjugated antibodies. Labelled cells were sorted on a FACSVantage flow cytometer (BD Biosciences, San Jose, CA, USA), and data analyzed using FlowJo software. CD45^+^ immune cells and CD31^+^ endothelial cells were excluded during the cell sorting procedure. Mammary gland lineage-specific gene expression (*Krt5, Krt18,* and *Pdgfr*) in the purified cell populations was checked by quantitative PCR (qPCR) as reported (Di-Cicco et al, 2015, Peuhu et al, 2017). RNA was reverse-transcribed with MMLV H(-) Point reverse transcriptase (Promega, Madison, WI, USA), and qPCR was performed by monitoring, in real time, the increase in fluorescence of the SYBR Green dye in a LightCycler^®^ 480 Real-Time PCR System (Roche Applied Science, Basel, Switzerland). The primers used for qPCR analysis (*Vimentin* and *Gapdh*) were purchased from SABiosciences/Qiagen (Hilden, Germany).

### Western blotting

The immunoblotting was done by using standard western blotting techniques and Odyssey LICOR imaging system.

### Cleared fat pad transplantation

Cleared fat pad transplantation was conducted as in (Peuhu et al, 2017). Briefly, mammary gland pieces (approx. 1 mm^3^) from 15 weeks old wt or *Vim-/-* female donor mice were transplanted under isoflurane anesthesia and analgesia (Temgesic, Rimadyl) to the fourth fat pad of 3 weeks old wt hosts (wt and *Vim-/-* transplants on each side of the host) after clearing the fat pad up to, and including, the lymph node. The removed part of the fat pad was fixed and stained to confirm complete removal of the recipient mouse mammary epithelium (Suppl. Fig.1). Growth of transplants was analyzed after 10 weeks by Carmine-alum staining of the fourth mammary gland whole mounts. For secondary transplantation, one mouse with wt and *Vim-/-* primary transplants in each fat pad was chosen randomly, both fat pads were excised, and an area close to the site of primary transplantation in each fat pad was cut into 5 pieces. These pieces were further transplanted in the fourth fat pads of 3 weeks old wt recipient mice (wt and *Vim-/-* transplants on each side of the host). Growth of transplants was analyzed after 15 weeks by Carmine-alum staining of the fourth mammary gland whole mounts. Growth take-on-rate was calculated from all the transplant samples.

### Statistical analysis

Sample size for the studies was chosen according to previous studies in the same area of research. A minimum of 3 mice was analysed for each genotype for comparison, and the data were assumed to follow a normal distribution. The GraphPad program and Student’s t-test (paired or unpaired, two-tailed) were used for all statistical analyses. Data are presented in column graphs with mean ± standard error of mean (SEM) and p-values. Individual data points per condition are shown when n≤15 and n-numbers are indicated in Figure legends.

## Author contributions

EP and JI contributed to the conception and design of the study. RV and AM conducted and analysed in vitro experiments. RV, EP and AW performed in vivo experiments and IHC analysis. EP conducted the cleared fat pad transplantations. EP wrote the manuscript and JI, RV and AW edited the manuscript. JI supervised the research.

## Acknowledgements

We would like to thank J. Jukkala, J. Siivonen, P. Laasola and M. Saari for instrumental technical assistance, Dr. H. Hamidi for editing of the manuscript, Dr. E. Närva, Dr. C. Guzman, and Dr. G. Jacquemet for critical reviewing of the manuscript, and Dr. M.-A. Deugnier and A. Di-Cicco for vimentin qPCR analysis. Prof. J. Eriksson is thanked for sharing the vimentin knockout mouse strain and Dr. F. Cheng for assistance in sample preparation. The Turku Centre for Biotechnology Cell Imaging Core, University of Turku Central Animal Laboratory, and Biocenter Finland are acknowledged for services, instrumentation and expertise.

## References

Asiedu MK, Beauchamp-Perez FD, Ingle JN, Behrens MD, Radisky DC, & Knutson KL (2014) AXL induces epithelial-to-mesenchymal transition and regulates the function of breast cancer stem cells. Oncogene 33: 1316–1324

Borgna S, Armellin M, di Gennaro A, Maestro R, & Santarosa M (2012) Mesenchymal traits are selected along with stem features in breast cancer cells grown as mammospheres. Cell Cycle 11: 4242–4251

Bouras T, Pal B, Vaillant F, Harburg G, Asselin-Labat ML, Oakes SR, Lindeman GJ, & Visvader JE (2008) Notch signaling regulates mammary stem cell function and luminal cell-fate commitment. Cell Stem Cell 3: 429–441

Brooks DL, Schwab LP, Krutilina R, Parke DN, Sethuraman A, Hoogewijs D, Schorg A, Gotwald L, Fan M, Wenger RH, & Seagroves TN (2016) ITGA6 is directly regulated by hypoxia-inducible factors and enriches for cancer stem cell activity and invasion in metastatic breast cancer models. Mol Cancer 15: 26-016-0510-x

Castillo-Lluva S, Hontecillas-Prieto L, Blanco-Gomez A, Del Mar Saez-Freire M, Garcia-Cenador B, Garcia-Criado J, Perez-Andres M, Orfao A, Canamero M, Mao JH, Gridley T, Castellanos-Martin A, & Perez-Losada J (2015) A new role of SNAI2 in postlactational involution of the mammary gland links it to luminal breast cancer development. Oncogene 34: 4777–4790

Cheng F, Shen Y, Mohanasundaram P, Lindstrom M, Ivaska J, Ny T, & Eriksson JE (2016) Vimentin coordinates fibroblast proliferation and keratinocyte differentiation in wound healing via TGF-beta-Slug signaling. Proc Natl Acad Sci U S A 113: E4320–7

Colucci-Guyon E, Gimenez Y Ribotta M, Maurice T, Babinet C, & Privat A (1999) Cerebellar defect and impaired motor coordination in mice lacking vimentin. Glia 25: 33–43

Colucci-Guyon E, Portier MM, Dunia I, Paulin D, Pournin S, & Babinet C (1994) Mice lacking vimentin develop and reproduce without an obvious phenotype. Cell 79: 679–694

Coulombe PA & Wong P (2004) Cytoplasmic intermediate filaments revealed as dynamic and multipurpose scaffolds. Nat Cell Biol 6: 699–706

Desgrosellier JS, Lesperance J, Seguin L, Gozo M, Kato S, Franovic A, Yebra M, Shattil SJ, & Cheresh DA (2014) Integrin alphavbeta3 drives slug activation and stemness in the pregnant and neoplastic mammary gland. Dev Cell 30: 295–308

Di-Cicco A, Petit V, Chiche A, Bresson L, Romagnoli M, Orian-Rousseau V, Vivanco M, Medina D, Faraldo MM, Glukhova MA, & Deugnier MA (2015) Paracrine Met signaling triggers epithelial-mesenchymal transition in mammary luminal progenitors, affecting their fate. Elife 4 doi: 10.7554/eLife.06104

Eckes B, Colucci-Guyon E, Smola H, Nodder S, Babinet C, Krieg T, & Martin P (2000) Impaired wound healing in embryonic and adult mice lacking vimentin. J Cell Sci 113 (Pt 13): 2455–2462

Green, MC, & Witham, BA (2009) The Jackson Laboratory Handbook on Genetically Standardized Mice.

Guo W, Keckesova Z, Donaher JL, Shibue T, Tischler V, Reinhardt F, Itzkovitz S, Noske A, Zurrer-Hardi U, Bell G, Tam WL, Mani SA, van Oudenaarden A, & Weinberg RA (2012) Slug and Sox9 cooperatively determine the mammary stem cell state. Cell 148: 1015–1028

Jackson HW, Waterhouse P, Sinha A, Kislinger T, Berman HK, & Khokha R (2015) Expansion of stem cells counteracts age-related mammary regression in compound Timp1/Timp3 null mice. Nat Cell Biol 17: 217–227

Kim J, Yang C, Kim EJ, Jang J, Kim SJ, Kang SM, Kim MG, Jung H, Park D, & Kim C (2016) Vimentin filaments regulate integrin-ligand interactions by binding to the cytoplasmic tail of integrin beta3. J Cell Sci 129: 2030–2042

Korsching E, Packeisen J, Liedtke C, Hungermann D, Wulfing P, van Diest PJ, Brandt B, Boecker W, & Buerger H (2005) The origin of vimentin expression in invasive breast cancer: epithelial-mesenchymal transition, myoepithelial histogenesis or histogenesis from progenitor cells with bilinear differentiation potential? J Pathol 206: 451–457

Kreis S, Schonfeld HJ, Melchior C, Steiner B, & Kieffer N (2005) The intermediate filament protein vimentin binds specifically to a recombinant integrin alpha2/beta1 cytoplasmic tail complex and co-localizes with native alpha2/beta1 in endothelial cell focal adhesions. Exp Cell Res 305: 110–121

Liu CY, Lin HH, Tang MJ, & Wang YK (2015) Vimentin contributes to epithelial-mesenchymal transition cancer cell mechanics by mediating cytoskeletal organization and focal adhesion maturation. Oncotarget 6: 15966–15983

Mailleux AA, Overholtzer M, Schmelzle T, Bouillet P, Strasser A, & Brugge JS (2007) BIM regulates apoptosis during mammary ductal morphogenesis, and its absence reveals alternative cell death mechanisms. Dev Cell 12: 221–234

Mani SA, Guo W, Liao MJ, Eaton EN, Ayyanan A, Zhou AY, Brooks M, Reinhard F, Zhang CC, Shipitsin M, Campbell LL, Polyak K, Brisken C, Yang J, & Weinberg RA (2008) The epithelialmesenchymal transition generates cells with properties of stem cells. Cell 133: 704–715

Mendez MG, Kojima S, & Goldman RD (2010) Vimentin induces changes in cell shape, motility, and adhesion during the epithelial to mesenchymal transition. FASEB J 24: 1838–1851

Nassour M, Idoux-Gillet Y, Selmi A, Come C, Faraldo ML, Deugnier MA, & Savagner P (2012) Slug controls stem/progenitor cell growth dynamics during mammary gland morphogenesis. PLoS One 7: e53498

Nieminen M, Henttinen T, Merinen M, Marttila-Ichihara F, Eriksson JE, & Jalkanen S (2006) Vimentin function in lymphocyte adhesion and transcellular migration. Nat Cell Biol 8: 156–162

Park D, Xiang AP, Mao FF, Zhang L, Di CG, Liu XM, Shao Y, Ma BF, Lee JH, Ha KS, Walton N, & Lahn BT (2010) Nestin is required for the proper self-renewal of neural stem cells. Stem Cells 28: 2162–2171

Peuhu E, Kaukonen R, Lerche M, Saari M, Guzman C, Rantakari P, De Franceschi N, Warri A, Georgiadou M, Jacquemet G, Mattila E, Virtakoivu R, Liu Y, Attieh Y, Silva KA, Betz T, Sundberg JP, Salmi M, Deugnier MA, Eliceiri KW et al (2017) SHARPIN regulates collagen architecture and ductal outgrowth in the developing mouse mammary gland. EMBO J 36: 165–182

Rahal RM, de Freitas-Junior R, Carlos da Cunha L, Moreira MA, Rosa VD, & Conde DM (2011) Mammary duct ectasia: an overview. Breast J 17: 694–695

Rangel MC, Bertolette D, Castro NP, Klauzinska M, Cuttitta F, & Salomon DS (2016) Developmental signaling pathways regulating mammary stem cells and contributing to the etiology of triple-negative breast cancer. Breast Cancer Res Treat 156: 211–226

Rios AC, Fu NY, Lindeman GJ, & Visvader JE (2014) In situ identification of bipotent stem cells in the mammary gland. Nature 506: 322–327

Shackleton M, Vaillant F, Simpson KJ, Stingl J, Smyth GK, Asselin-Labat ML, Wu L, Lindeman GJ, & Visvader JE (2006) Generation of a functional mammary gland from a single stem cell. Nature 439: 84–88

Soady KJ, Kendrick H, Gao Q, Tutt A, Zvelebil M, Ordonez LD, Quist J, Tan DW, Isacke CM, Grigoriadis A, & Smalley MJ (2015) Mouse mammary stem cells express prognostic markers for triple-negative breast cancer. Breast Cancer Res 17: 31-015-0539-6

Taddei I, Deugnier MA, Faraldo MM, Petit V, Bouvard D, Medina D, Fassler R, Thiery JP, & Glukhova MA (2008) Beta1 integrin deletion from the basal compartment of the mammary epithelium affects stem cells. Nat Cell Biol 10: 716–722

Tiede B & Kang Y (2011) From milk to malignancy: the role of mammary stem cells in development, pregnancy and breast cancer. Cell Res 21: 245–257

Vaillant F, Asselin-Labat ML, Shackleton M, Forrest NC, Lindeman GJ, & Visvader JE (2008) The mammary progenitor marker CD61/beta3 integrin identifies cancer stem cells in mouse models of mammary tumorigenesis. Cancer Res 68: 7711–7717

Van Keymeulen A, Rocha AS, Ousset M, Beck B, Bouvencourt G, Rock J, Sharma N, Dekoninck S, & Blanpain C (2011) Distinct stem cells contribute to mammary gland development and maintenance. Nature 479: 189–193

Virtakoivu R, Mai A, Mattila E, De Franceschi N, Imanishi SY, Corthals G, Kaukonen R, Saari M, Cheng F, Torvaldson E, Kosma VM, Mannermaa A, Muharram G, Gilles C, Eriksson J, Soini Y, Lorens JB, & Ivaska J (2015) Vimentin-ERK Signaling Uncouples Slug Gene Regulatory Function. Cancer Res 75: 2349–2362

Visvader JE & Stingl J (2014) Mammary stem cells and the differentiation hierarchy: current status and perspectives. Genes Dev 28: 1143–1158

Vuoriluoto K, Haugen H, Kiviluoto S, Mpindi JP, Nevo J, Gjerdrum C, Tiron C, Lorens JB, & Ivaska J (2011) Vimentin regulates EMT induction by Slug and oncogenic H-Ras and migration by governing Axl expression in breast cancer. Oncogene 30: 1436–1448

Wang D, Cai C, Dong X, Yu QC, Zhang XO, Yang L, & Zeng YA (2015) Identification of multipotent mammary stem cells by protein C receptor expression. Nature 517: 81–84

Yamashita N, Tokunaga E, Kitao H, Hisamatsu Y, Taketani K, Akiyoshi S, Okada S, Aishima S, Morita M, & Maehara Y (2013) Vimentin as a poor prognostic factor for triple-negative breast cancer. J Cancer Res Clin Oncol 139: 739–746

Ye X, Tam WL, Shibue T, Kaygusuz Y, Reinhardt F, Ng Eaton E, & Weinberg RA (2015) Distinct EMT programs control normal mammary stem cells and tumour-initiating cells. Nature 525: 256–260

Zelenko Z, Gallagher EJ, Tobin-Hess A, Belardi V, Rostoker R, Blank J, Dina Y, & LeRoith D (2017) Silencing vimentin expression decreases pulmonary metastases in a pre-diabetic mouse model of mammary tumor progression. Oncogene 36: 1394–1403

Zeng YA & Nusse R (2010) Wnt proteins are self-renewal factors for mammary stem cells and promote their long-term expansion in culture. Cell Stem Cell 6: 568–577

